# Semantic segmentation of microscopic neuroanatomical data by combining topological priors with encoder-decoder deep networks

**DOI:** 10.1101/2020.02.18.955237

**Authors:** Samik Banerjee, Lucas Magee, Dingkang Wang, Xu Li, Bingxing Huo, Jaik-ishan Jayakumar, Katie Matho, Adam Lin, Keerthi Ram, Mohanasankar Sivaprakasam, Josh Huang, Yusu Wang, Partha P. Mitra

## Abstract

Understanding of neuronal circuitry at cellular resolution within the brain has relied on tract tracing methods which involve careful observation and interpretation by experienced neuroscientists. With recent developments in imaging and digitization, this approach is no longer feasible with the large scale (terabyte to petabyte range) images. Machine learning based techniques, using deep networks, provide an efficient alternative to the problem. However, these methods rely on very large volumes of annotated images for training and have error rates that are too high for scientific data analysis, and thus requires a significant volume of human-in-the-loop proofreading. Here we introduce a hybrid architecture combining prior structure in the form of topological data analysis methods, based on discrete Morse theory, with the best-in-class deep-net architectures for the neuronal connectivity analysis. We show significant performance gains using our hybrid architecture on detection of topological structure (e.g. connectivity of neuronal processes and local intensity maxima on axons corresponding to synaptic swellings) with precision/recall close to 90% compared with human observers. We have adapted our architecture to a high performance pipeline capable of semantic segmentation of light microscopic whole-brain image data into a hierarchy of neuronal compartments. We expect that the hybrid architecture incorporating discrete Morse techniques into deep nets will generalize to other data domains.

## Introduction

Understanding the morphology and the connectivity of neurons is an important step in determining the neural circuitry of the brain. A primary method used for this purpose in larger vertebrate brains involves sparsely labelling individual neurons or groups of neurons using neuronal tracer injections. These methods have been the gold standard in neuroanatomical circuit tracing studies because they directly visualize neurons with high SNR and no inferential process is needed. These data sets have traditionally been painstakingly examined under a microscope by expert neuroanatomists, or more recently using digital microscopy, but still employing human labor intensive methods requiring close and time-consuming interactions with a human expert. Modern advances in optical imaging have allowed the digitization of whole-brain data sets, either 2D image stacks or 3D image volumes, with subcellular resolution^1–3^. These imaging techniques have been incorporated into integrated neurohistology pipelines to generate large-scale, high-resolution data sets in a high-throughput fashion^4–6^.The resulting multi-Terabyte to Petabyte scale data sets are not possible to study using purely manual detection and quantification. Thus there is a pressing need for efficient automated algorithms to achieve this goal, with high precision and recall suitable for scientific data analysis.

Machine Learning techniques have facilitated the processing and analysis of large neuroanatomical data sets. In particular, computer algorithms for detecting neuronal fibers from various images with high accuracy have become increasingly important in automating computational characterization of neuronal morphology and circuitry. Previous works include reconstructions of neuronal morphology ^7^, neuronal data analysis with an emphasis on reconstructing neurons from EM data cubes^8^; digital reconstruction of the 3D morphology of neurons from image stacks^9^. Recent advancements include a large-scale automated server-based biomedical-image analysis in FARSIGHT^10^, a free and open-source toolkit of image analysis methods for quantitative studies of complex and dynamic tissue micro-environments. The BigNeuron project ^(i)^ is a community effort to advance the state of the art of single-neuron reconstruction^11^. There is a also a large literature on the related but broader field of semantic segmentation of biomedical image data, including histopathological data, which we briefly review below.

In this manuscript, we develop a computational framework for a systematic treatment of the semantic segmentation problem for neuroanatomical image data. The basis of our framework is a hierarchical set of semantic categories suitable for neuroanatomical images (Figure 1). A key component of this framework is a new method for automated segmentation of neurites (axons and dendrites), combining topological data analysis, based on Discrete Morse (DM) theory (Figure Methods 1), together with encoder-decoder Deep Net architectures to detect and quantify neuronal processes and boutons in whole brain histology sections. We call this method DM++ (Figure 2). Usage of the Morse-theoretic topological prior allows us to retain the fitting flexibility of deep-learning frameworks, while at the same time incorporating non-trivial prior structure. The resulting method shows both quantitative and qualitative improvements over the existing state of the art encoder-decoder networks for the same task (Table 1). Although neurite detection is a key step in the semantic segmentation task, there are also other components to the framework (Figure 3, Figure Supplementary 3) to detect other relevant compartments such as somata, or to further sub-categorize detected neurites (Figure 5). We have deployed our methodology into a semi-automated data analysis pipeline including human proofreaders to obtain high quality semantic segmentation of the neuroanatomical image data (Figure Supplementary 2), suitable for further neuroscientific analysis.

**Table 1:** shows the Precision, Recall, F1 values for the different tasks. The unsupervised technique for Process Detection involves intensity-based thresholding with hard-coded parameters for threshold and morphological operations. Cell Detection Results are shown for the WSI fluorescent images (MBA) and WSI Bright Field images (BFI) compared to a baseline technique (SegNET). The (I) on the side of the MaskRCNN indicates that we omit Injection Region while reporting our metrics. In computing these metrics we utilize a 5-pixel neighborhood of a detected point to judge detects. The values in bold indicate the best performance.

| Model | Precision | Recall | F1 |
| --- | --- | --- | --- |
| **Process Detection - STP** |  |  |  |
| Unsupervised | 0.61 | 0.63 | 0.62 |
| UNET | 0.85 | 0.88 | 0.86 |
| ALBU | 0.90 | 0.88 | 0.89 |
| DM++ | **0.92** | **0.93** | **0.92** |
| **Process Detection - MBA** |  |  |  |
| Unsupervised | 0.42 | 0.51 | 0.46 |
| UNET | 0.67 | 0.73 | 0.70 |
| ALBU | 0.79 | 0.82 | 0.81 |
| DM++ | **0.83** | **0.84** | **0.84** |
| **Process Detection - BFI** |  |  |  |
| Unsupervised | 0.59 | 0.61 | 0.60 |
| UNET | 0.73 | 0.75 | 0.74 |
| ALBU | 0.79 | 0.80 | 0.80 |
| DM++ | **0.81** | **0.83** | **0.82** |
| **Cell Detection - BFI** |  |  |  |
| SegNet | 0.79 | 0.74 | 0.76 |
| MaskRCNN(I) | **0.85** | **0.89** | **0.87** |
| **Cell Detection - MBA** |  |  |  |
| SegNet | 0.72 | 0.75 | 0.73 |
| MaskRCNN | 0.81 | 0.82 | 0.82 |
| MaskRCNN(I) | **0.85** | **0.90** | **0.87** |
| **Passing Axon Detection - MBA** |  |  |  |
| SegNet | 0.72 | 0.76 | 0.74 |
| MaskRCNN | **0.89** | **0.93** | **0.91** |
| **Axon Arbor Detection - MBA** |  |  |  |
| SegNet | 0.72 | 0.75 | 0.74 |
| MaskRCNN | **0.95** | **0.97** | **0.96** |
| **Dendrite Detection - MBA** |  |  |  |
| SegNet | 0.67 | 0.71 | 0.69 |
| MaskRCNN | **0.79** | **0.86** | **0.83** |
| **Bouton Detection - MBA** |  |  |  |
| Discrete Morse | 0.76 | 0.31 | 0.44 |

**Figure 1:**
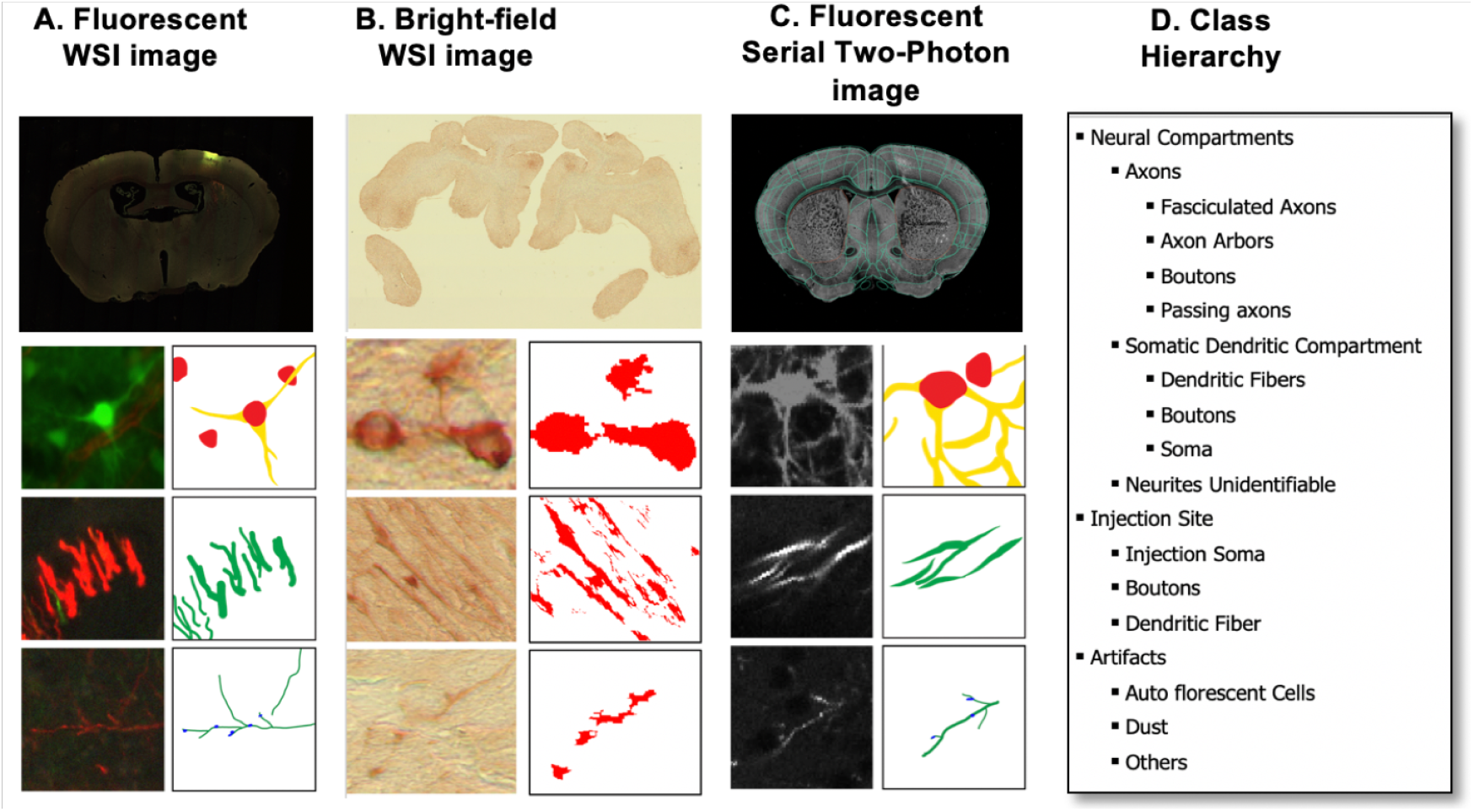
Three prevalent modalities of light-microscopic data acquisition where we have applied our algorithm: (**1A**) Colored fluorescent Whole Slide Images (WSI) of brain sections (mouse) acquired using digital slide scanners, (**1B**) Bright Field images of immunohistochemically brain sections (marmoset) and (**1C**) monochrome Serial Two Photon Tomographic images (STPT) (mouse). The top panels show the whole section images and below are the neuronal soma, neuronal processes and putative terminating axon arbors with putative synaptic boutons. **1D** shows the class hierarchy that we devised to categorize different image components (objects of interests) that we targeted for semantic segmentation.

**Figure 2:**
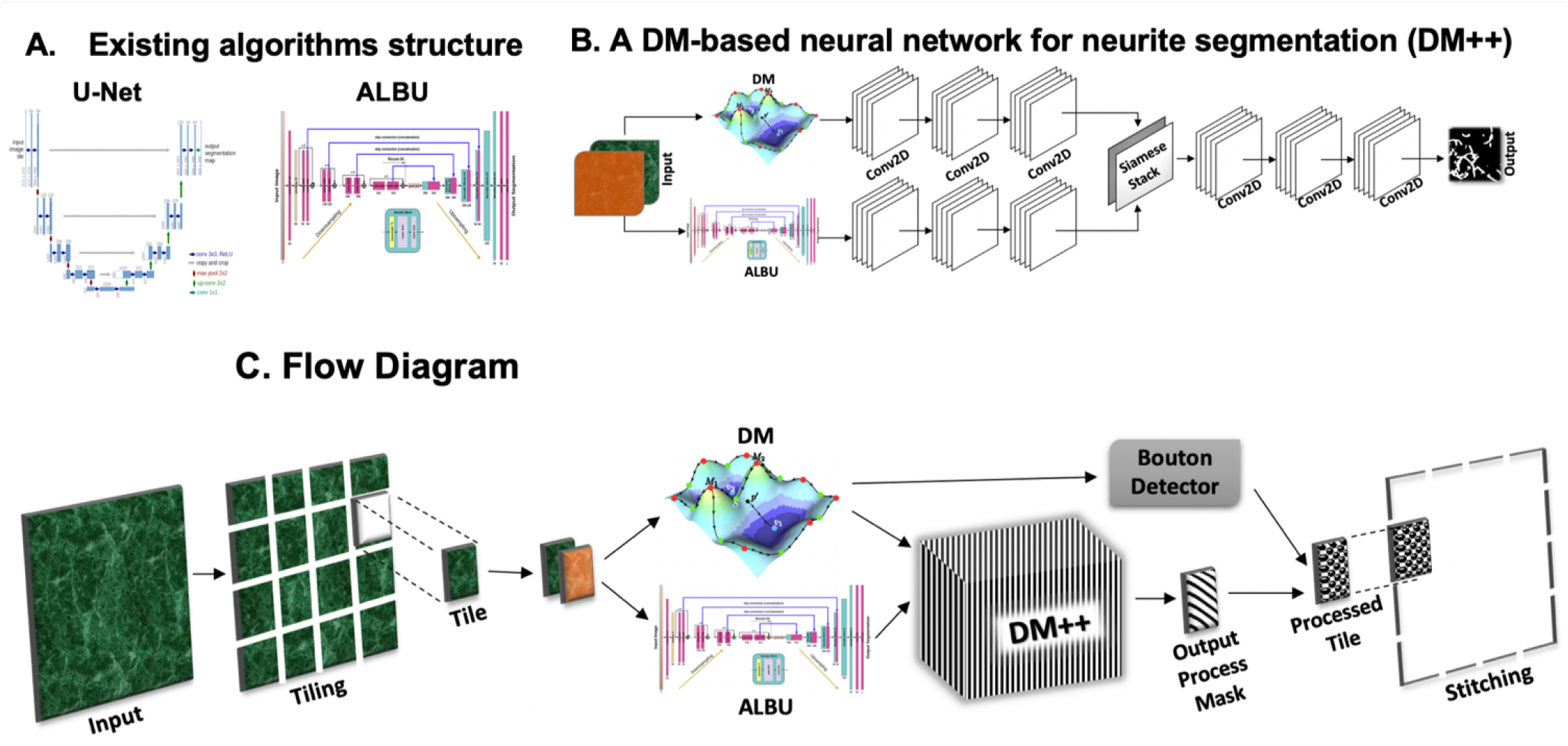
Existing CNN architectures suitable for neurite segmentation, U-Net (**2A**-left) and ALBU (**2A**-right) illustrated along with our proposed architecture of DM++ (**2B**). The DM++ architecture concatenates a “Topological path” and a “DCNN path” using a Siamese Stacking layer, followed by a final common CNN stack. (**2C**) shows the combined Process and Bouton Detection Pipeline.

**Figure 3:**
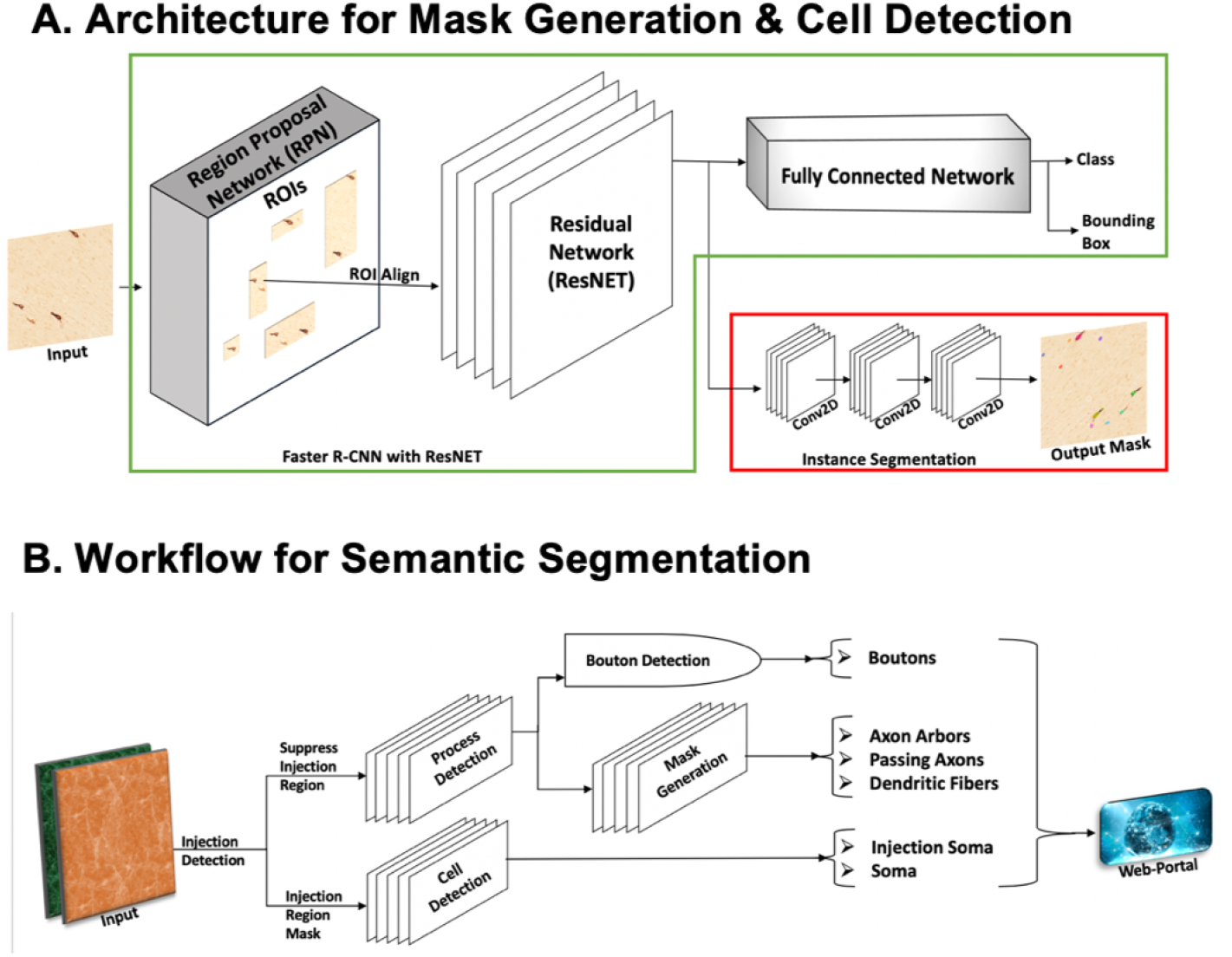
**(3A)** The figure shows the architecture of the Mask-RCNN network, which we use for Cell Detection as well as for mask generation for different process sub-classes (*e.g.*, passing vs arborized axons). **(3B)** Semantic segmentation for tracer-injected brains is preceded by an Injection Detection step. This is needed since the injection region has a different signal dynamic range than the projection regions. The injection region is suppressed for the Process Detection task, and is also used to separate “injection region somata” from distal somata. The Process Detection output (manually proof-read) acts as a mask for discarding the other background pixels while generating masks for process sub-classes.

## Background

Automated computer vision algorithms have been used extensively for the delineation of anatomical structures and other regions of interest in specific radiologica1 and histopathological images. These image segmentation algorithms, play an important role in numerous biomedical-imaging applications, such as the quantification of tissue volumes^12^, diagnosis^13^, localization of pathology^14^, study of anatomical structure^15^, treatment planning^16^, and computer-integrated surgery^17^.

Prior to the popularization of Deep CNN methods^18–20^, several machine learning and image processing techniques were available for medical image segmentation. These include amplitude segmentation based on histogram features^21^, the region based segmentation^22^, and the graph-cut approach^23^. However, semantic segmentation approaches that utilize DL have become popular in recent years in the field of medical image segmentation, lesion detection, and localization^24^. DCNN architectures have generally provided state-of-the-art performance for image classification ^25^, segmentation^26^, detection and tracking ^27^, and captioning ^28^ in standardized datasets. Efficient optimization techniques are available for training DCNN models^25^. However, in most cases, models are explored and evaluated on classification tasks in large-scale datasets like ImageNet^25^, where the outputs of the classification tasks are a single label or probability value for a given image. Alternatively, smaller fully-connected convolutional neural networks (FCN)^29^ and Segnet^30^ have been used successfully for semantic image segmentation tasks. State-of-the-art DCNN architectures for image segmentation tasks like U-Net^31^ and ALBU^32^ (Figure 2A) employ an encoder-decoder architecture. While these networks are effective they employ double the number of model parameters thus raising resource issues.

Deep Learning techniques rely on very large volumes of annotated training data and may in some sense be training-data interpolation techniques^33^. To address the lack of large annotated corpora for new tasks, data transformation or augmentation techniques^31, 34, 35^ including data whitening, rotation, translation, and scaling have been applied to increase the number of labeled samples available. The class imbalance problem is often solved by using tile based approaches rather than entire image sets^36^. However, these kinds of methodological fixes do not ultimately address the weakness of DL techniques in lacking domain-specific priors, such as the topological prior that can capture a better connectivity of the neuronal architecture in the brain for our study.

To incorporate topological prior structure suitable to neurites, we employ Discrete Morse Graph Reconstruction (Figure Methods 1), a method that has been developed and studied thoroughly over the past several years^37–40^. This algorithm combines persistent homology ^41^ and Discrete Morse Theory^42^ to extract underlying graph structures from density fields. The algorithm considers the global structure of the data rather than looking strictly at local information and can handle many negative qualities seen in imperfect data, such as noise and non-uniform sampling. The method has been used in many applications, such as reconstructing road networks from GPS traces^40^ and extracting filament structures from simulated dark matter density fields^43^.

### Our Approach

In this paper, we propose a comprehensive framework for the semantic segmentation tasks for light-microscopic neuroanatomical data and compare with state of the art approaches. Gigapixel sized images are handled by breaking up into tiles of manageable size. The problem of class imbalance, when the pixel-based approach adapted for the tiles is applied to the entire image, is overcome by assigning a label for the image backgrounds and a target class for the foreground during semantic segmentation. In addition, two methods including binary cross-entropy loss and dice similarity were used for efficient training of classification and segmentation tasks^34, 35^.

To detect neuronal processes, we propose a hybrid CNN based architecture DM++ (Figure 2B) which incorporates a topological prior using discrete Morse theory to give additional weight to quasi-one dimensional connected structures such as neurites. DCNNs do not have any built in mechanisms that can exploit the topological connectivity of neurites. For example, variations in the data acquisition process may cause some parts of a neurite to be less intensely labelled than other parts. This “signal gap” may pose challenges for a DCNN which might suppress the label on the weakly labelled part of a neurite (refer to the leftmost tile in the middle row for each modality in Figure 4). Nevertheless, prior knowledge that neurites form connected branches of tree-shaped neurons allow a human observer to easily trace through such regions of low intensity. We employ the Discrete Morse technique to incorporate such global connectivity information, as shown in the rightmost tile in the middle row for each modality in Figure 4.

**Figure 4:**
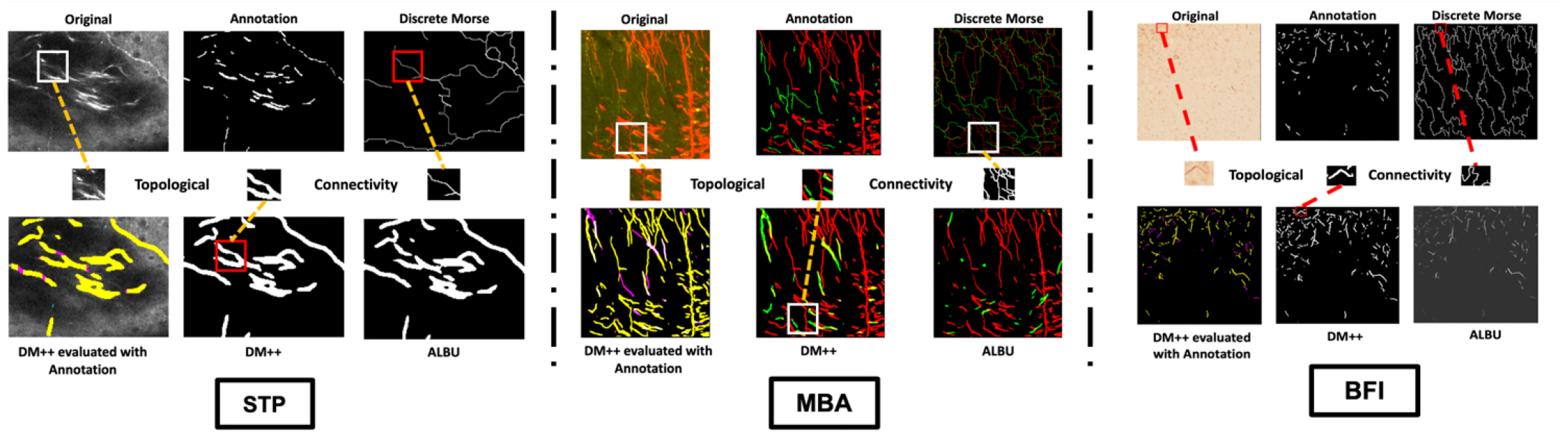
Illustrations of results from the process detection pipeline are shown here for single-color STP images **(3 leftmost columns)**, 3-color fluorescent WSI images from the Mouse Brain Architecture project data set (MBA) **(3 middle columns)**, and the Brightfield WSI images **(3 right most columns)**. The top left row shows the original image, human annotation of the processes, and the discrete Morse output. The bottom left shows evaluation of DM++ output compared with the human annotation, with true positives (TP) in yellow and “false positives” (FP) in red. The FP connect the human-annotated neurite fragments, and examination of the original image data shows low intensity signal connecting the annotated fragments. The final panel in the bottom left row is the ALBU output. In each of the three modalities, the middle row depicts a small patch from the image, where the intensity captured by the digital scanners used for imaging these sections is significantly low, but the connectivity can be observed in the high resolution image by naked human eye, when inspected meticulously (leftmost patch in the middle row). As evident in the detections by ALBU, a state-of-the-art CNN technique, they are generally missed while we generate the mask. These faint connections are well captured by Discrete Morse as evident in the patch from the same location in the output of the Discrete Morse reconstruction algorithm (rightmost patch in the middle row). With the introduction of the topological prior the patch from the same location of our output preserves the connectivity (central patch in the middle row). These patches demonstrate that the ALBU output misses faint connections that are picked up by the DM++ algorithm. The bottom right row shows the evaluation of DM++ output compared to manual annotations with TP (yellow), FP (magenta) and FN (cyan). In the MBA modality, the top right row shows red and green process fragments corresponding to neurites labelled with red and green fluorescent protein corresponding to anterograde AAV tracer injections, human annotations (double labelled process fragments are labelled green), and discrete morse output. Evaluations, DM++ and ALBU outputs for the colored MBA image data are shown next, followed by the similar structure in the BFI dataset. The rectangular box indicates the location for each image from where the exemplar patch is extracted and the dotted lines establishes the correspondences of each path to the middle row of results for better visibility.

We tested the proposed DM++ algorithm on three different types of image modalities used to image sections of the brain. Examples of the brain section images are shown in the top panels of columns A-C in figure 1, in each case from brains in which neuronal tracer injections have been used to visualize a subset of neurons and their processes. Figure 1(A) shows colored (fluorescent) Whole Slide Images (WSI). Figure 1(B) shows the Bright Field Images (BFI) of a brain section with an immunohistological stain. Both these images were captured using a digital slide scanner (0.46*μ*m/pixel). Figure 1(C) shows monochrome images captured using Serial Two Photon Tomography (STPT). The second row (Column 1 A-C) shows examples of labelled neuronal cell bodies (soma) in these different modalities. The next third row shows neuronal processes labelled by red Fluorescent protein (Column 1 A), immunohistochemistry using the ABC-DAB process (Column 1 B) and high intensity eGFP label from the STPT data set (Column 1 C). The bottom row (Column 1 A-C) show examples of putatively terminating neuronal axons with the synaptic swellings/boutons. The cartoons in the second, third and fourth row (Column 1 A-C) beside each image tiles (the right subcolumn) shows masks corresponding to the object categories of interest in each of the tiles shown in the left subcolumn. The semantic category hierarchy is detailed in the Column 1 D.

In the following sections, we first elaborate the pipeline for process detection, followed by the the rest of the semantic category tree. Subsequently we present results, followed by discussion and conclusions.

#### Neuronal Process detection: DM++ Architecture

The DCNN component of the DM++ pipeline is based on the ALBU network ^32^ which is in turn based on U-Net ^31^. Figure 2(A) illustrates U-net and ALBU. 2(C) shows a diagrammatic representation of our proposed architecture DM++ for process detection. A more extensive description can be found in the Methods Section.

Briefly, the DM++ network accepts an image containing relevant objects of interest (e.g. processes) as input, tiles the image, processes these tiles using three processing paths forming a Y-shaped network, and outputs a binary mask indicating the detected neuronal processes. Figure 2(B) shows the three paths, (i) A “topological path” for detecting connected neural processes in the tile, (ii) a “DCNN path” (ALBU) for detecting the processes (iii) the final “common path” for combining the DCNN and topological paths.

### Semantic Segmentation

Brain section images not only contain information about one object of interest (processes as detected by the DM++ pipeline) but also other objects such as dendrites, soma. Axonal processes themselves can show heterogeneity depending on their terminating nature. To account for these detections, we have incorporated the mask-RCNN architecture^26^. This modification enables us to determine the exact pixels belonging to a sub-process or somata. The detections for the sub-categories of processes use the process detection masks generated by the DM++ algorithm. The architecture of Mask-RCNN, developed on Faster-RCNN with an instance segmentation module, takes in the output neuronal mask from the DM++, to detect the axon arbors, passing axons and dendrites (for detailed discussion on the Mask-RCNN architecture, please refer to the Methods section). In contrary, the cell detection is performed by the semantic segmentation pipeline independent of the neuronal processes, though the boutons are detected as a sub-category using the output vertices of the graph from the DM reconstruction algorithm within the DM++ pipeline. A brief discussion on the bouton detection is given in the methods section. A pictorial representation of the the overall flow diagram for the semantic segmentation methodology determined for the neuronal segmentation with different class labels and the cell detection has been depicted in figure 3 (B), while figure 3 (A) shows the architecture for the Mask-RCNN used to generate masks for each of the sub-categories of the neuronal processes as well as cell detection.

## Results

The outputs from the proposed algorithms of neuronal process detection and semantic segmentation discussed above have been evaluated both qualitatively and quantitatively. This section provides a discussion of these evaluations, when compared with other state-of-the-art techniques. The proposed methodology shows improved performance metrics, and in addition also shows the ability to better detect connected process fragments. Finally, we also discuss the time complexities of the proposed pipeline, in terms of training and testing, followed by a discussion on the improvement in annotation time induced by our auto-detects.

### Results: Neuronal Process detection

The results are presented in table 1. The table shows the Precision, Recall and F1-measure for the different modalities. We have compared DM++ with an Unsupervised, UNet and ALBU for neuronal process detection. Each of these techniques are discussed in the Methods section. As evident from the sub-tables, our proposed technique outperforms the state of the art architectures across performance measures.

Figure 4 shows example results for different image modalities. Of particular interest is portions of neuronal processes connected by regions of low signal intensity, where DM++ is able to connect the processes through these low-intensity regions, but ALBU does not do so. In each of the three modalities, the middle row depicts a small patch from the image, where the intensity captured by the digital scanners used for imaging these sections is significantly low, but the connectivity can be detected in the high resolution image by visual inspection (refer to the leftmost patch in the middle row of results for each modality).

In each of these modalities, we also show comparisons with the manually annotated data (demarcated as “Annotation” in the figure) with TP (yellow), FP (magenta) and FN (cyan). The rectangular box indicates the location for each image from where the exemplar patch is extracted and the dotted lines establishes the correspondences of each path to the middle row of results for precise inspection. This kind of preservation of connectivity was evident throughout the sections used for testing. This is consistent with the expectation that the topological prior would improve the detection of connected processes.

### Semantic Segmentation

The semantic segmentation pipeline illustrated in figure 3 was used for detecting the sub-neuronal classes and somata. Performance metrics are presented in table 1. The numbers in bold indicate the results from our overall architecture, compared with a state-of-the-art encoder-decoder based CNN architecture, SegNet^30^ (details in Methods section).

Figure 5 illustrates each of the classes discussed in figure 1. Results for Cell Detection are shown in figure 5(A), which includes the MBA and the BFI dataset. The different sub-compartments of neuronal processes are displayed for the MBA dataset in the subsequent figures. Figures 5(B-D) illustrate the results of the semantic segmentation pipeline for different sub-categories of neuronal processes. The output from the DM++ algorithm is provided as input to the MaskRCNN network, as shown in the top tile. The output mask from the segmentation pipeline is shown just below, while the bottom row shows the performance of the MaskRCNN when compared to the manually annotated ground truth. Precision-Recall-F1 scores are calculated and reported in table 1.

**Figure 5:**
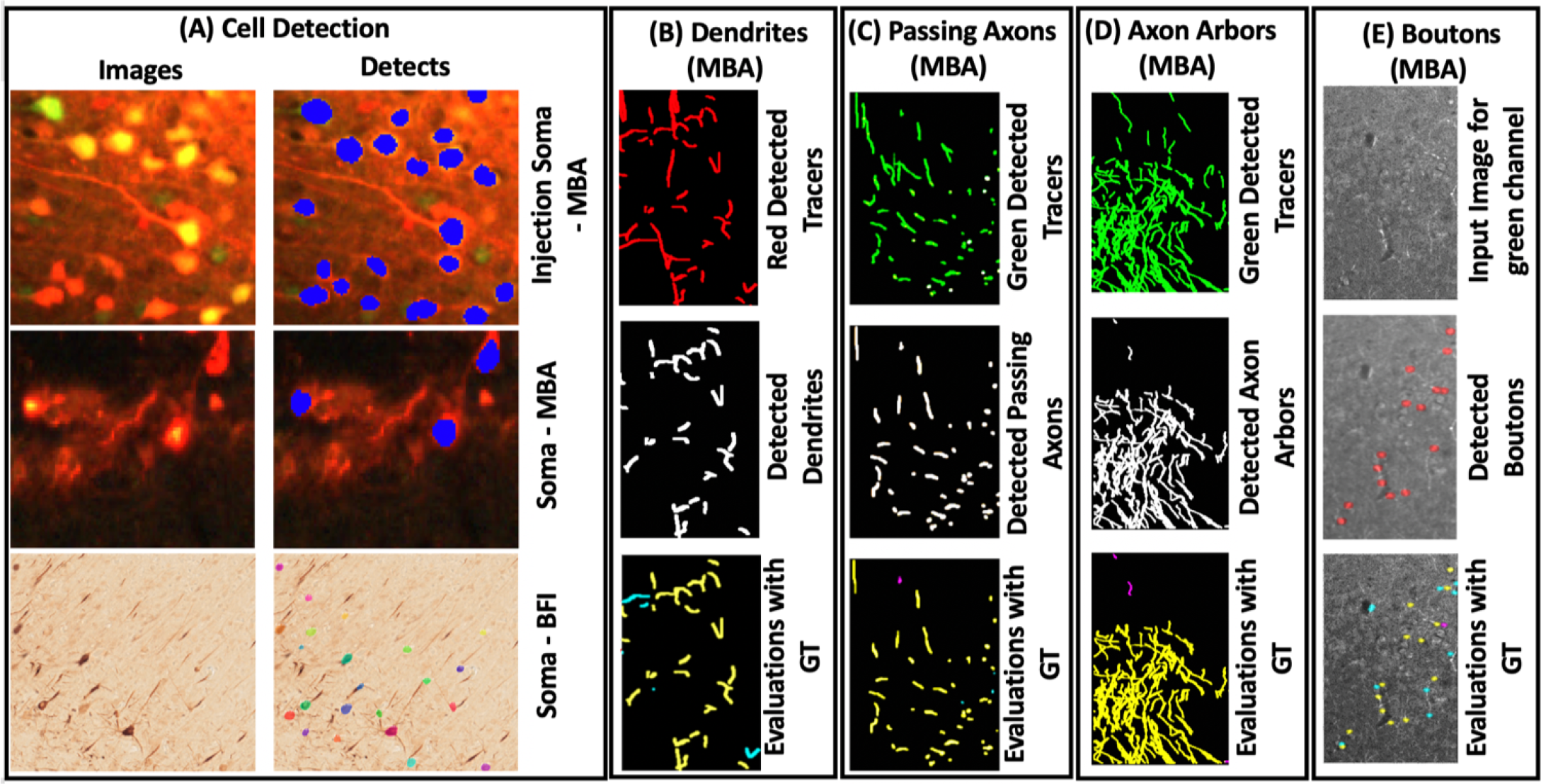
Semantic Segmentation results: **(A)** the results for cell detection for the two modalities along with the injection soma detection for the MBA dataset. Soma segmentations are shown to the immediate right of the original image shown in the left column. The right column shows the detected cells in a different color, for the corresponding left patch. The top row shows a region where the tracer is injected (“Injection Region”). The detected cells in this region are labelled “Injection Soma”. The bottom two rows show detections outside this injection region for the two modalities, Fluorescent WSI images (MBA data) and Brightfield WSI images. **(B)** The results for the semantic category (“dendrites”) for MBA data, a sub-category of processes. **(C)** The results for the semantic category (“passing axons”) for MBA data, a sub-category of processes. **(D)** The results for the semantic category (“axon arbors”) of the MBA data. The top tile in each of these semantic categories shows the detected processes from the Process Detection Pipeline for the green/red tracer. The middle tile illustrates the resulting semantic segmentation mask, while the bottom tile shows the comparison of our result with the Groundtruth (GT) annotation, depicting the TP (yellow), FP (magenta) and FN (cyan) for the input tile. **(E)** The results for bouton detection on the MBA dataset.

As an additional usage of the Discrete Morse approach, we detect putative “boutons” on axons by finding intensity maxima along the detected proesses. The resulting “bouton” detections are shown in 5(E). The first image is an input tile, the second image shows what our method outputs as putative boutons, and the third image identifies whether these are true positives or false positives (compared with expert judgment), as well as where false negatives occur. Modified precision, recall, and F1 scores for three tiles that were manually annotated are included in 1. These modified scores were obtained with using a radius of 5, meaning a bouton labeled by our method would be labeled as a true positive if the nearest true bouton was within a neighborhood of 5 pixels (2.5*μ*).

### Computational metrics

We trained the DM++ network for 500 epochs with the training data. The training time was 200 seconds per epoch. For process detections, the algorithm took 4 minutes per mouse brain section (each section is a 24*K* × 24*K* image with 0.46*μ*m/pixel resolution) with processing times of 12% pre-processing, 15% neuronal pathway, 48% Topological pathway, 15% combined pathway and 10% post-processing, on a dual GPU (NVIDIA GT×2080 Ti) machine. The topological pathway was run in parallel on a 36 core CPU with 40 parallel threads. We used the pyTorch library for the python code for the neuronal pathway and Keras with TensorFlow backend for the common pathway.

We used Keras with Tensorflow backend for the MaskRCNN network for cell detection and mask-generation. The Cell Detection MaskRCNN network was trained for 200 epochs, with a training time of ~1000 seconds per epoch. The detects took an average of 3 minutes per section for the cell (soma) detections on the mouse brain dataset. The architecture used for mask generation follows the one-vs-all strategy and approximately takes 3 minutes on an average for the three classes combined into an unified framework. There is an significant improvement in the proof-reading times for the manual annotations. Based on the report of an experienced neuroanatomist, the proof-reading time for the process detection is no more than 25% compared to annotating the processes de-novo. A similar trend was seen for cell-detection where we can reduced the annotation time by ~70%. In each case, proofreading was done with equally high quality criteria as de-novo manual annotation.

## Discussion and Conclusions

We demonstrate significant performance improvements for an image segmentation task over state-of-the-art encoder-decoder networks through the incorporation of prior topological structure using methods from Morse theory (Table 1 and Figures 4 & 5). We show that such priors may be incorporated using an intuitively plausible Y-shaped (“Siamese”) network architecture we entitle DM++. We further incorporate this process-detection network as an important component in an integrated framework for semantic segmentation of light microscopic neuroanatomical image data. While we present significant performance improvements over state of the art, we are not yet at the point where the automation is at a sufficiently high quality level for scientific purposes, and a human-in-the loop component is required (although the trained networks lead to a large reduction of the required human labor). Our post-hoc analysis of the data shows that the DM++ algorithm cannot handle abrupt changes in the dynamic intensity range within the data set, so that training datasets containing a full spectrum of intensity dynamic range is required. We found however that our networks incrementally increase performance based feedback from manual curation, which can be used to repeatedly fine-tune the network to improve generalization. Despite these residual shortcomings, our new algorithm (DM++) incorporating topological priors, using discrete Morse theory in combination with DCNNs, significantly outperforms existing techniques for the detection and segmentation of neuronal objects of interests in light-microscopic neuroanatomical data.

## Code and Dataset Availability

The code and documentation will be available on Github on article publication.

## Acknowledgements

The authors gratefully acknowledge IHC Brightfield Imaged data from Dr Peter Strick at U Pitt, and wish to thank Jaimi Nagashima and Mitsutoshi Hanada also at U Pitt for annotating these images. We acknowledge the effort from annotators at the Center for Computational Brain Research at IIT Madras for the bulk of the data annotation and proofreading for this project. This work was supported by the NIH (EB022899, MH114824, MH114821, NS107466, AT010414), the Crick-Clay Professorship (Cold Spring Harbor Laboratory), the Mathers Charitable Foundation, and H. N. Mahabala Chair (IIT Madras). Work at Ohio State University was in addition partly supported by the NSF under grants CCF-1740761, RI-1815697, and DMS-1547357.

## Author Contributions

The idea of using topological priors in the pipeline was conceptualized by Y.W. and P.M. Algorithmic design and development was performed by S.B and L.M. Proof reading assistance and neuroanatomical expertise for neuroanatomical ground truth data was provided by J.J. and K.M. Data preparation, including quality control and acquisition were performed by B.H, J.J. and K.M. under the supervision of J.H. and P.M. Evaluation of the algorithm was conducted by S.B., L.M., X.L. Data preparation, including design of an online proofreading interface and hosting was done by A.L., M.S. and K.R. S.B., L.M., J.J. and P.M. prepared and edited the paper

## Methods

This section describes methods used in this study including the fundamentals of the algorithm design, the architecture, training, testing and evaluation of our technique when compared to the existing conventional methods. We have compared our pipeline to existing techniques include an unsupervised process detection technique, U-Net and ALBU for process detection and SegNet for semantic segmentation, which are also discussed at the end of the Methods.

### Discrete Morse Graph Reconstruction

DM++ takes masks of the Discrete Morse Graph Reconstruction output as input. To generate this, we first need a density function over the domain to feed into the Discrete Morse framework. Image intensity in the color channel relevant to the tracer serves as the density function after applying a Gaussian filter to smooth out the image. We can interpret our density function as the higher the pixel value, the more likely it is to be part of true neuron signal.

Next we input the density function into the Discrete Morse Graph reconstruction framework. If one were to imagine the density function graphed on top of the domain, the Discrete Morse framework outputs the “mountain ridges” of the function, connecting the maxima through intervening saddle points. These ridges are made up of “flow lines”^(ii)^ between the maxima and saddle points of the function. The reason for selecting these is that the function values are locally maximal along the ridges. Thus if we were to move slight off of a ridge, the probability of being part of the neuronal process decreases.

#### Noise Removal

Each ridge is assigned a persistence value^41^ which can be interpreted as an importance score. The noise-removal framework adopts a persistence threshold and filters out ridges below that threshold. We provide a low threshold value in order to remove ridges caused by noise while minimizing the number of ridges removed that actually mark true neuron signal.

Given the Morse output, we then create a mask for the domain. The mask is grayscale - where each path in the Discrete Morse output is assigned a constant value. The output of this stage is a simple grayscale image which can then be used at the next step of the DM++ algorithm. The higher the intensity along the path, the higher the gray scale value. After the persistence refined Discrete Morse graph reconstruction step, this mask, along with the original imaging data, is an input to the process detection pipeline (explained under the subsequent subsection DM++).

### DM++

The combination of the topological component with a DCNN convolutional network is aimed to capturing the connected nature of the neuronal processes. The “Siamese” architecture of the DM++ network takes the outputs from the discrete Morse algorithm and the ALBU architecture and combines them using a “Siamese” stacking into two channels. The common path emerging out of the Siamese stack combines features of the convolutional network along with the topological priors of the Discrete Morse.

#### Architecture

The architecture of DM++ consists of four components: the topological path, the ALBU path, the Siamese stack and the common path. The DCNN paths in this network consist of two-dimensional convolutional layers with ReLU activations, except the final layers in each which have Sigmoid activations. A dropout layer of 50% was introduced in each DCNN stack for regularization purposes.

#### Inputs

The inputs to the DM++ architecture are the likelihood maps obtained from the discrete Morse algorithm and the ALBU models. The input images are assumed to be single channel images with the resolution of 512 × 512 pixels with varying pixel resolutions across different modalities of the images. The input image is passed through the discrete Morse algorithm to obtain a likelihood image containing the connected graphs with their edge weights as their likelihood. The ALBU model also produces output for the likelihood maps generated for segmentation of the processes.

#### Outputs

The output of DM++ produces a likelihood map of the processes in 512 × 512 pixel tiles. The outputs are in the range [0, 255], with a higher value indicating greater confidence that a pixel belongs to a process present in the data. These likelihood maps were thresholded at a predefined threshold (STP: 100, MBA: 75 and BFI: 120) to generate a mask for the process in the section.

### Pipeline for Process Detection

The overall flow of the brain image data from the input to output is denoted as a pipeline for neuronal process detection. The pipeline for Process Detection is illustrated in figure 2(B), which was designed for two-dimensional process detection for a tracer injection in the brain, run section-wise for each brain. Each of these sections are first broken down into tiles of 512 × 512 pixels. These tiles are then individually passed to the discrete Morse algorithm and the trained ALBU models. The gray-scale maps generated out of the discrete Morse algorithm are fed into the topological pathway of the DM++ and the segmentation likelihood predicted by the ALBU models is fed to the neuronal segmentation pathway. These priors pass through these pathways and then merge into a dual stack (can be thought of as a two-channel image). This stack is further propagated through the common pathway deeper into the network to captures the high-level features of the neuronal process based on both the priors. The trained DM++ network outputs a likelihood map for the process which is further thresholded to a binary mask for identifying the processes. This mask is also used to localize the detection of boutons, and the sub-categories of the neuronal processes discussed later in the semantic segmentation pipeline. These output tiles are eventually stitched together to form a cumulative mask for the whole section. The discrete Morse algorithm also outputs the peaks in the images as the vertices in the graph, which are seen in along the processes as the high-intensity peaks, identified as boutons or synaptic swellings on the neurons.

**Figure Methods 1:**
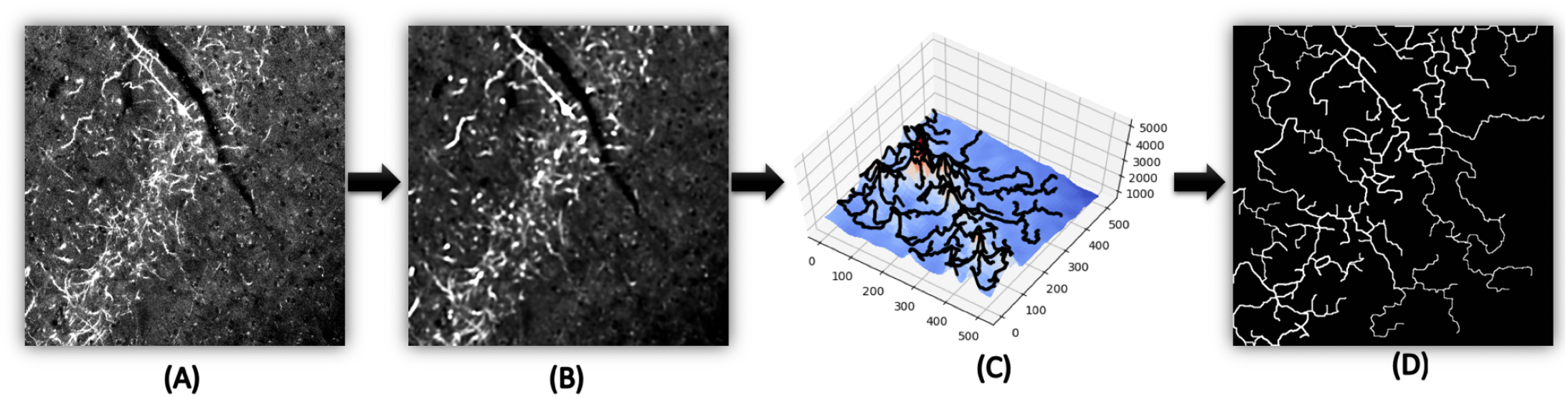
The Discrete Morse algorithm is given an input image **(A)**. A Gaussian filter is applied to the image **(B)** - a density function is defined at the pixels. Then the algorithm extracts the ridges of the function across the domain **(C)** - these ridges form the 1-stable manifold. Finally, each path in the 1-stable manifold is assigned an grayscale value based on intensity along the path, and a grayscale mask is outputted in **(D)**.

#### Variations in the pipeline for Process Detection

The pipeline has some variations across the different imaging modalities. The input to the pipeline is the raw image of a section from a brain for each data modality. The input is not the same across the different modalities, as seen in figure 1(A-C). The images from the Serial Two-Photon images are 16-bit gray scale images, while the images from the brains in Mouse Brain Architecture project are three-channel 12-bit images and the images from the brightfield WSI images were three-channel 8-bit images. The tracer injections in the STP and the Rabies data were of a single color but the MBA data had two injection colors.

The input to DM++ are single-color images. So, for the STP date the pipeline works on the raw data, while for the Brightfield images the 8-bit color data were converted into a single grayscale image before passing into the pipeline. The MBA data is treated channel-wise. Since the injections are green and red only, we assume that the green and red channels are separete likelihood images for the two tracers, and pass them individually through the pipeline, finally combining their outputs for the whole section. The BFI dataset is also treated as a likelihood image of intensities for the DM reconstruction but ALBU is performed on the three-channel color images.

#### Ground-truth Annotation

Manual annotation was used to generate training labels our supervised algorithms. The training set for the STP data consists of a coarse annotation of a whole brain (267 sections) and four section of high-quality precise annotations from another brain. The training data for the MBA images included four sections each from two brains with different labels marked as described in our class-hierarchy. The manually annotated BFI dataset consisted of 72 tiles of 1024 × 1024 pixels from a single section, which was divided for training and testing.

#### Preparation of the Data for training and testing

The manually labelled sections were broken down into tiles of size (512 × 12) pixels. Since the processes are really sparse in the whole brain, the fully annotated STP was randomly tiled to incorporate tiles with varying densities of annotations ranging down to no annotations. Hence, we had a training set with 5749 tiles from the fully annotated STP brain and two high-quality annotated sections entirely tiled to obtain another 750 tiles. The testing for the STP data was done on another two sections fully tiled of the precise annotation data. The eight sections from the two brains in the MBA data were carefully annotated. We used 7 sections for training, resulting in ~ 2000 tiles and testing was done on a section with ~ 500 tiles. The BFI data had 180 tiles in the training set and 108 tiles in the testing set. The total number of tiles for training and testing has been tabulated in table Supplementary 1, which gives an overview of the limited data available for training and testing our supervised networks.

#### Evaluation Metrics

The metrics used throughout the manuscript are Precision (P), Recall (R), and the F1-measure. For the evaluation of these metrics we determine the True Positives (TP), False Positives (FP), True Negatives (TN) and False Negatives (FN) (refer figure 4, bottom row leftmost tile for each modality). Every detected pixel was evaluated with a disk of radius of 5 pixels. if an annotated groundtruth existed within thid disk, the pixel was considered as TP, otherwise it is considered as FP. If an annotation did not have detected pixel within a 5-pixel radius, then we considered that as TN. Utilizing all these values, Precision and recall were computed as 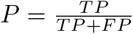 and 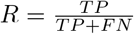, while the F1-score was calculated as harmonic mean of precision and recall, i.e., 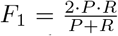. Also, we did an iterative refinement of annotations, as shown in the supplementary figure Supplementary 2, based on the FP detected by the algorithms on a first pass to double check for the human errors in the annotation. Each of our automatic detections were proof-read to ensure the authenticity of the automatic detects as shown in supplementary figure Supplementary 3.

**Table Supplementary 1:** The number of manually annotated tiles used for training and testing the different semantic categories in the different modalities of the brain. These manually annotated tiles are of size 512 × 512 pixels. They are used to train the different pipelines in the brain, with data augmentation, to cope with the limited availability of the training data. These tiles account for the tiles where the annotators have at least annotated any of the neuronal structures.

| Image modality | Neuronal classes | Tiles for training | Tiles for testing |
| --- | --- | --- | --- |
| MBA | Processes | 2000 | 494 |
| STPT | Processes | 6449 | 704 |
| BFI | Processes | 180 | 108 |
| MBA | Soma Detection | 26 | 9 |
| BFI | Soma Detection | 48 | 24 |
| MBA | Dendrites | 23 | 8 |
| MBA | Passing Axons | 22 | 6 |
| MBA | Axon Arbors | 22 | 6 |

### Bouton Detection

The Discrete Morse Graph Reconstruction algorithm was combined with the finalized process detection outputs to detect putative synaptic boutons. A persistent homology ^41^ computation was performed within the Discrete Morse Graph Reconstruction. Boutons appear in the image data as small balls with relatively high intensity along neuronal structures. This means that they will be vertices along the Morse skeleton graph with relatively high persistence. Thus to perform bouton detection, we took the vertices of the Morse skeleton output with persistence greater than a user provided threshold. Then from these points, we only include those that fell within the final process detection mask. Portions of the Morse skeleton graph lie outside of true process signal, so such a filter is needed to avoid labeling boutons outside of true process signals.

### Pipeline for Semantic Segmentation

The semantic segmentation architecture included a MaskRCNN network (see figure 3(A)) which performed two segmentation tasks: (i) cell detection, to identify the somata in the injection region as well as in other parts of the brain; and (ii) classification of the different process classes (axon arbors, fasciculated axons, and passing axons). For the semantic segmentation of the before-mentioned classes, the Process Detection is taken as a pre-processing step to mask out the background. MaskRCNN was employed on this pre-processed image to determine all the classes denoted in figure 1(D).

#### Architecture of MaskRCNN

MaskRCNN architecture (figure 3(A)) included a FasterRCNN network, which has two outputs for each candidate object, a class label, and a bounding-box offset. To this network, we add a third branch that outputs the object mask — which is a binary mask that indicates the pixels where the object is in the bounding box. The additional mask output is distinct from the class and box outputs, requiring extraction of much finer spatial layout of an object. To do this MaskRCNN uses the Fully Convolution Network (FCN). FasterRCNN is a good algorithm that is used for object detection. FasterRCNN consists of two stages. The first stage called a Region Proposal Network (RPN), which proposes candidate object bounding boxes. The second stage extracts features using RoIPool from each candidate box and performs classification and bounding-box regression. The features used by both stages can be shared for inference. So, in short, we can say that MaskRCNN combines the two networks — FasterRCNN and FCN - into one joint architecture. The loss function for the model is the combined loss in doing classification, generating a bounding box and generating the mask.

#### Soma Detection

The soma detection strategy is two-fold: (i) soma detection within the injection region, termed as “Injection soma” and (ii) soma detection outside the injection region. The injection region is either determined previously by their higher intensities for the different colors in the tracers or they are manually demarcated by an expert. At the injection region, we perform a contrast enhancement and intensity normalization using histogram equalization. All of the soma detected within this region are demarcated as the injection soma, while the others are referred as projection-region soma. Both of these are detected as masks by the MaskRCNN architecture. These masks are determined as individual connected components in the brain slices and the centroids of these connected components are treated as cell-centers. The soma detection algorithm was also applied to all the sections for the MBA (please refer to table Supplementary 1 for the size of the training and testing set); both injection and non-injection sections. For the BFI dataset, limited data availability (please refer to table Supplementary 1 for the size of the training and testing set) allowed us only to categorize the data as soma (where we assume to be outside any injection-like region). The evaluation of these cell centers are also based on the same evaluation technique described earlier.

#### Semantic Detection

The semantic detection deals with the second layer of the hierarchy after Process Detection. The processes consist mostly of axons and dendrites, but for the semantic detections we are targeting the different intrinsic properties of these processes - *viz.* the axon arbors and the passing axons. Similarly we sub-categorize dendritic processes.

On a training set of four manually annotated sections, we trained the MaskRCNN network individually on each of these classes using one-vs-other classification strategy, with data augmentation. The results for the sub-categories from the axons and dendrites are masked using the process detection mask generated earlier. This semantic segmentation process was carried on the MBA data. The overall pipeline of the semantic segmentation is described in figure 3(B) where the process detection acts as one of the important modules within the semantic segmentation pipeline. The evaluations for the semantic detections also follow the same procedure as discussed earlier for the process detection.

First, we detect the dendrites. The training is done on four sections. The rest is classified as axons. The manual annotation had 31 tiles of size 512 × 512 pixels to contain data. Out of which 8 tiles have been separated out for the testing and rest for training. Secondly, on the detected axons, we train our MaskRCNN on the passing axons. The rest are classified as axon arbors. In total, from the 4 sections we had 28 tiles with axonal data. The model trained on dendrites throws out the dendrites from the process detection masks. Six of these axonal tiles have been used for testing while the rest are used for training the MaskRCNN (please refer to table Supplementary 1 for the size of the training and testing set). The quantitative values are reported in table 1.

### Techniques under comparison

#### Unsupervised Process Detection

We assumed that the intensity of neuronal processes are much higher compared to the background and formulated an unsupervised algorithm based on the classical image processing techniques and morphological operations on the input image. The input image is first thresholded at a particular intensity (~ 0.4 for STPT data, ~ 0.3 channel-wise for the MBA data), which suppresses the background. Edge detection is then performed on the thresholded image using the “Canny” edge detector^44^, to extract the long connected edges in the images. The longer connected components from the blurred image are extracted from the images using binary morphological operations image opening and closing followed by area opening, which eliminates the small components or the artifacts in the image. The remaining connected components are classified as the process or the output of our unsupervised technique.

#### UNet

UNet^31^ is a generic deep-learning solution for frequently occurring quantification tasks such as cell detection and shape measurements in biomedical image data. The main idea is to supplement a contracting network by successive layers, where pooling operations are replaced by upsampling operators, thus increasing the resolution of the output. A successive convolutional layer can then be used to assemble a precise output based on this information. One important modification in U-Net is that there are a large number of feature channels in the upsampling part, which allow the network to propagate context information to higher resolution layers. As a consequence, the expansive path is more or less symmetric to the contracting part, and yields a u-shaped architecture. The contracting path is a typical convolutional network that consists of repeated application of convolutions, each followed by a rectified linear unit (ReLU) and a max pooling operation. During the contraction, the spatial information is reduced while feature information is increased. The expansive pathway combines the feature and spatial information through a sequence of up-convolutions and concatenations with high-resolution features from the contracting path. There are many applications of U-Net in biomedical image segmentation, such as brain image segmentation (“BRATS”^45^) and liver image segmentation^46^.

#### ALBU

We used the ALBU ^32^ network as a strong baseline for process detection. The ALBU network has a U-Net-like architecture. It is composed of an encoder and a decoder with skip connections. The encoder is a pre-trained 34-layer residual neural network (ResNet-34 ^47^), and the decoder consists of sequential decoder blocks. Each decoder block starts with an upsample layer followed by two convolution layers^32^ (See figure 2(A)). The ALBU network has significant advantages over U-Net and other U-Net based architectures. It was originally applied on satellite image segmentation tasks and was the winner of the road detection challenge from SpaceNet ^(iii)^.

#### SegNet

SegNet^30^, is designed to be an efficient architecture for pixel-wise semantic segmentation. This overall architecture consists of an encoder network, a corresponding decoder network followed by a pixel-wise classification layer. It is also significantly smaller in the number of trainable parameters than other competing architectures and can be trained end-to-end using stochastic gradient descent. The authors^30^ present a real-time online demo of road scene segmentation into 11 classes of interest for autonomous driving to demonstrate the effectiveness of SegNet. We adopted SegNet as a baseline for comparison with our pipeline for semantic segmentation, since the results produced by SegNet were not adequate. We then moved to MaskRCNN as our core for the pipeline as it outperformed SegNet and produced better results.

---

**Figure Supplementary 1:**
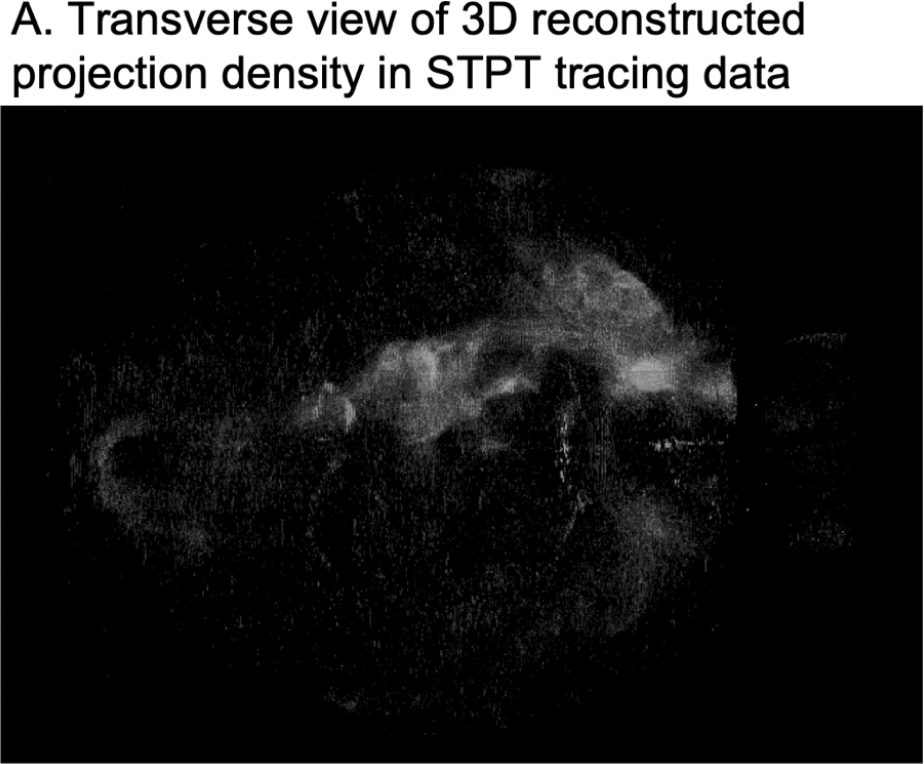
Processes detection and 3D reconstruction.

## Supplementary Materials

### Image modalities

In mammalian brains, tract tracing using tracer injections is considered the gold standard for studying long range connectivity. The data sets included in our analysis come from mesoscale connectivity mapping studies using tracer injections, with brains digitized using a variety of imaging techniques. We utilized data from three different imaging modalities, including two whole-slide imaging data sets with in-plane resolution of 0.46*μ* (brightfield, three-color, 8-bits per color images as well as fluorescent three-color, 12-bits per color images), and one single-color serial two-photon tomographic data set (16 bit images, in-plane resolution 1*μ*).

### Future work on summarizing tracer injections

The future scope of work includes the design of a combined pipeline for all the datasets - robust to different modalities. The data summarization into a 3D volume for the visualization of the whole brain is also a step to explore. Preliminary results of the 3D summarization of the process detection is illustrated in figure Supplementary 1. A study on such summarization has been conducted in the accompanying manuscript ^48^.

### Improvement in Annotation and trained models due to manual curation of results

Manual annotation is a time-consuming process and depends on the annotator to judge and annotate the processes individually. This annotation technique is subjective based on the perception of an individual annotator. Thus, building on an initial small set of annotations to train our model, the iterative refinement of the annotation is an important and necessary step. We demonstrate the results of this iterative process in figure Supplementary 2, where we term this process of incremental improvement by manual intervention as a ‘Feedback Network’, as shown in subfigure A. An examples from the STPT dataset, shown in subfigure B, depicts how the annotation is refined using our Feedback Network.

**Figure Supplementary 2:**
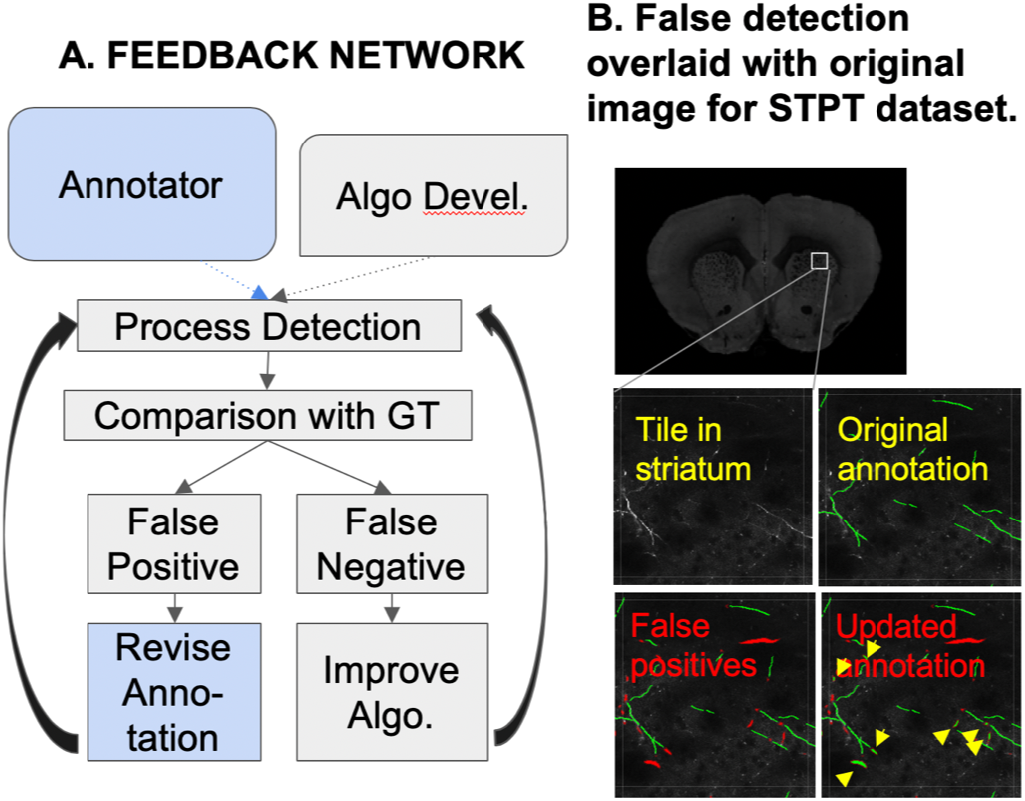
**(A)** Illustration for the framework of iterative proofreading of the automatic annotations. **(B)** An example from the STP dataset showing the correction after first stage of automatic process detection.

### Overall Pipeline

The overall pipeline depicts an evolving strategy towards the fully-automatic neuroanatomical data analysis. The yellow blocks in the workflow in figure Supplementary 3 signifies the manual curation step expected to be operative till our models achieves high performance over all brain images across modalities. In the future, these blocks can serve as quality control steps to ensure validity of the predictions.

**Figure Supplementary 3:**
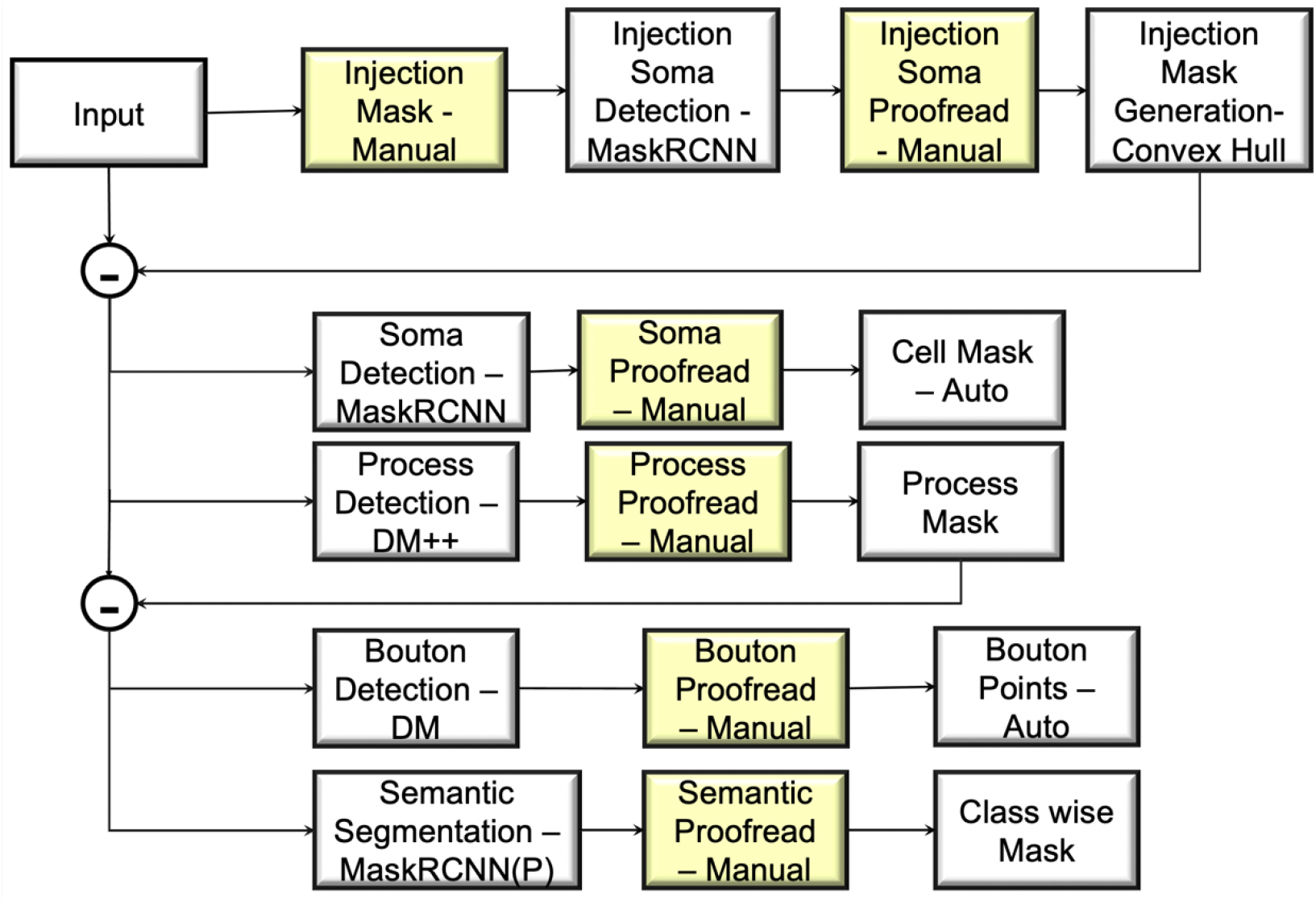
The overall workflow for the whole semantic segmentation. The yellow squares indicates the point of manual proof-reading of the automatically generated results. The (P) indicates that this module uses process detection mask to eradicate the other pixels.

## Footnotes

(i) http://bigneuron.org/

(ii) The relevant flow lines (more formally integral lines) are curves whose local tangents follow the direction of steepest descent, i.e, the gradient direction, of the density function.

(iii) https://github.com/SpaceNetChallenge/RoadDetector

## References

1. Bohland, J. W. et al. A proposal for a coordinated effort for the determination of brainwide neuroanatomical connectivity in model organisms at a mesoscopic scale. PLoS computational biology 5(2009).

2. Dodt, H.-U. et al. Ultramicroscopy: three-dimensional visualization of neuronal networks in the whole mouse brain. Nature methods 4, 331–336 (2007).

3. Ragan, T. et al. Serial two-photon tomography for automated ex vivo mouse brain imaging. Nature methods 9, 255 (2012).

4. Oh, S. W. et al. A mesoscale connectome of the mouse brain. Nature 508, 207–214 (2014).

5. Pinskiy, V. et al. High-throughput method of whole-brain sectioning, using the tape-transfer technique. PloS one 10(2015).

6. Lin, M. K. et al. A high-throughput neurohistological pipeline for brain-wide mesoscale connectivity mapping of the common marmoset. Elife 8, e40042 (2019).

7. Halavi, M., Hamilton, K. A., Parekh, R. & Ascoli, G. Digital reconstructions of neuronal morphology: three decades of research trends. Frontiers in neuroscience 6, 49 (2012).

8. Helmstaedter, M. & Mitra, P. P. Computational methods and challenges for large-scale circuit mapping. Current opinion in neurobiology 22, 162–169 (2012).

9. Peng, H., Meijering, E. & Ascoli, G. A. From diadem to bigneuron (2015).

10. Rey-Villamizar, N. et al. Large-scale automated image analysis for computational profiling of brain tissue surrounding implanted neuroprosthetic devices using python. Frontiers in neuroinformatics 8, 39 (2014).

11. Peng, H. et al. Bigneuron: large-scale 3d neuron reconstruction from optical microscopy images. Neuron 87, 252–256 (2015).

12. Lawrie, S. M. & Abukmeil, S. S. Brain abnormality in schizophrenia: a systematic and quantitative review of volumetric magnetic resonance imaging studies. The British Journal of Psychiatry 172, 110–120 (1998).

13. Taylor, R. H., Lavealle, S., Burdea, G. C. & Mosges, R. Computer-integrated surgery: technology and clinical applications (Mit Press, 1995).

14. Zijdenbos, A. P. & Dawant, B. M. Brain segmentation and white matter lesion detection in mr images. Critical reviews in biomedical engineering 22, 401–465 (1994).

15. Worth, A. J., Makris, N., Caviness Jr, V. S. & Kennedy, D. N. Neuroanatomical segmentation in mri: technological objectives. International Journal of Pattern Recognition and Artificial Intelligence 11, 1161–1187 (1997).

16. Khoo, V. S. et al. Magnetic resonance imaging (mri): considerations and applications in radiotherapy treatment planning. Radiotherapy and Oncology 42, 1–15 (1997).

17. Grimson, W. E. L. et al. Utilizing segmented mri data in image-guided surgery. International Journal of Pattern Recognition and Artificial Intelligence 11, 1367–1397 (1997).

18. LeCun, Y. Yoshua bengio, and geoffrey hinton. Deep learning. nature 521, 436–444 (2015).

19. LeCun, Y., Bottou, L., Bengio, Y. & Haffner, P. Gradient-based learning applied to document recognition. Proceedings of the IEEE 86, 2278–2324 (1998).

20. Fukushima, K. Neocognitron: A self-organizing neural network model for a mechanism of pattern recognition unaffected by shift in position. Biological cybernetics 36, 193–202 (1980).

21. Ramesh, N., Yoo, J.-H. & Sethi, I. Thresholding based on histogram approximation. IEE Proceedings-Vision, Image and Signal Processing 142, 271–279 (1995).

22. Sharma, N. & Ray, A. K. Computer aided segmentation of medical images based on hybridized approach of edge and region based techniques. In Proc. Int. Conf. Math. Biol (2006).

23. Boykov, Y. Y. & Jolly, M.-P. Interactive graph cuts for optimal boundary & region segmentation of objects in nd images. In Proceedings eighth IEEE international conference on computer vision. ICCV 2001, vol. 1, 105–112 (IEEE, 2001).

24. Litjens, G. et al. A survey on deep learning in medical image analysis. Medical image analysis 42, 60–88 (2017).

25. Krizhevsky, A., Sutskever, I. & Hinton, G. E. Imagenet classification with deep convolutional neural networks. In Advances in neural information processing systems, 1097–1105 (2012).

26. He, K., Gkioxari, G., Dollár, P. & Girshick, R. Mask r-cnn. In Proceedings of the IEEE international conference on computer vision, 2961–2969 (2017).

27. Redmon, J., Divvala, S., Girshick, R. & Farhadi, A. You only look once: Unified, real-time object detection. In Proceedings of the IEEE conference on computer vision and pattern recognition, 779–788 (2016).

28. Vinyals, O., Toshev, A., Bengio, S. & Erhan, D. Show and tell: Lessons learned from the 2015 mscoco image captioning challenge. IEEE transactions on pattern analysis and machine intelligence 39, 652–663 (2016).

29. Sabour, S., Frosst, N. & Hinton, G. E. Dynamic routing between capsules. In Advances in neural information processing systems, 3856–3866 (2017).

30. Badrinarayanan, V., Kendall, A. & Cipolla, R. Segnet: A deep convolutional encoder-decoder architecture for image segmentation. IEEE transactions on pattern analysis and machine intelligence 39, 2481–2495 (2017).

31. Ronneberger, O., Fischer, P. & Brox, T. U-net: Convolutional networks for biomedical image segmentation. In International Conference on Medical image computing and computer-assisted intervention, 234–241 (Springer, 2015).

32. Buslaev, A., Seferbekov, S. S., Iglovikov, V. & Shvets, A. Fully convolutional network for automatic road extraction from satellite imagery. In CVPR Workshops, 207–210 (2018).

33. Belkin, M., Hsu, D. J. & Mitra, P. Overfitting or perfect fitting? risk bounds for classification and regression rules that interpolate. In Advances in neural information processing systems, 2300–2311 (2018).

34. Çiçek, Ö., Abdulkadir, A., Lienkamp, S. S., Brox, T. & Ronneberger, O. 3d u-net: learning dense volumetric segmentation from sparse annotation. In International conference on medical image computing and computer-assisted intervention, 424–432 (Springer, 2016).

35. Milletari, F., Navab, N. & Ahmadi, S.-A. V-net: Fully convolutional neural networks for volumetric medical image segmentation. In 2016 Fourth International Conference on 3D Vision (3DV), 565–571 (IEEE, 2016).

36. Johnson, J. M. & Khoshgoftaar, T. M. Survey on deep learning with class imbalance. Journal of Big Data 6, 27 (2019).

37. Delgado-Friedrichs, O., Robins, V. & Sheppard, A. Skeletonization and partitioning of digital images using discrete morse theory. IEEE transactions on pattern analysis and machine intelligence 37, 654–666 (2014).

38. Gyulassy, A., Bremer, P.-T., Hamann, B. & Pascucci, V. A practical approach to morse-smale complex computation: Scalability and generality. IEEE Transactions on Visualization and Computer Graphics 14, 1619–1626 (2008).

39. Robins, V., Wood, P. J. & Sheppard, A. P. Theory and algorithms for constructing discrete morse complexes from grayscale digital images. IEEE Transactions on pattern analysis and machine intelligence 33, 1646–1658 (2011).

40. Dey, T. K., Wang, J. & Wang, Y. Road network reconstruction from satellite images with machine learning supported by topological methods. In Proceedings of the 27th ACM SIGSPATIAL International Conference on Advances in Geographic Information Systems, 520–523 (2019).

41. Edelsbrunner, H. & Harer, J. Persistent homology-a survey. Contemporary mathematics 453, 257–282 (2008).

42. Forman, R. A user’s guide to discrete morse theory. Sém. Lothar. Combin 48, 35pp (2002).

43. Sousbie, T., Pichon, C., Colombi, S., Novikov, D. & Pogosyan, D. The 3d skeleton: tracing the filamentary structure of the universe. Monthly Notices of the Royal Astronomical Society 383, 1655–1670 (2008).

44. Ding, L. & Goshtasby, A. On the canny edge detector. Pattern Recognition 34, 721–725 (2001).

45. Menze, B. H. et al. The multimodal brain tumor image segmentation bench-mark (brats). IEEE transactions on medical imaging 34, 1993–2024 (2014).

46. Kainmüller, D., Lange, T. & Lamecker, H. Shape constrained automatic segmentation of the liver based on a heuristic intensity model. In Proc. MICCAI Workshop 3D Segmentation in the Clinic: A Grand Challenge, 109–116 (2007).

47. He, J. et al. Deep reinforcement learning with a natural language action space. arXiv preprint arXiv:1511.04636 (2015).

48. Dingkang, W. et al. Detection and skeletonization of tracer injections using topological methods. In preparation.

